# Differential vaginal *Lactobacillus* species metabolism of glucose, L and D-lactate by ^13^C-nuclear magnetic resonance spectroscopy

**DOI:** 10.1101/2020.03.10.985580

**Authors:** Emmanuel Amabebe, Dilly O. Anumba, Steven Reynolds

## Abstract

**Introduction:** Cervicovaginal dysbiosis can lead to infection-associated spontaneous preterm birth.

**Objective:** To determine whether vaginal *Lactobacillus* species, *L. crispatus* and *L. jensenii*, differentially metabolise glucose, L- and/or D-lactate to propagate their survival/dominance.

**Methods:** Bacteria were incubated anaerobically for 24h at 37°C, with ^13^C_u_-glucose, ^13^C_3_-D-lactate or ^13^C_3_-L-lactate (singularly or combined) for 24h. ^13^C-spectra were acquired using a 9.4T NMR spectrometer.

**Results:** *L. crispatus* and *L. jensenii* (n=6 each) metabolised ^13^C-glucose to ^13^C-lactate and ^13^C-acetate. *L. jensenii* converted more ^13^C_3_-D- or ^13^C_3_-L-lactate to ^13^C-acetate than *L. crispatus*, p<0.001.

**Conclusion:** Conversion of glucose and lactate to acetate by *L. jensenii* compared to *L. crispatus*, suggests a possibly important pathomechanism of dysbiosis and infection-associated spontaneous preterm birth.

## Introduction

In conjunction with the host vaginal habitat, the vaginal bacteria produce unique metabolic by-products. In a healthy vagina, lactobacilli are the predominant species and have been linked to increased likelihood of term delivery (Amabebe and Anumba, 2018; Stafford *et al*., 2017). The four major vaginal lactobacilli (*L. crispatus*, *L. jensenii*, *L. iners* and *L. gasseri*) differentially produce L- and D-lactic acid that lowers the pH of the cervicovaginal space, creating unfavourable conditions for other invading species. D-lactic acid is believed to be more potent than L-lactic acid in relation to protection against colonisation by potentially pathogenic organisms within the vagina and the accompanying inflammatory response (Witkin *et al*., 2013). Anaerobes are also endogenous to the vagina and are associated with infection and preterm birth (PTB, i.e. delivery <37 weeks of gestation) especially when lactobacilli are deficient (Amabebe and Anumba, 2018).

Metabolism by cells can be tracked by ^13^C-Nuclear Magnetic Resonance (NMR) to examine specific pathways (Buescher *et al*., 2015). The use of ^13^C labelled substrates means that they are metabolised identically to those found naturally, with the cell viability maintained throughout the experiment. Additionally, strategic placement of the ^13^C label (Buescher *et al*., 2015) allows the activity of single or multiple metabolic pathways to be identified, even if the end product is the same (Bruntz *et al*., 2017).

Although, lactobacilli are known to thrive in the acidic condition of the vagina (pH < 4.5), the mechanism underpinning how lactobacilli, and other anaerobes, interact with lactic acid remains unresolved (Amabebe and Anumba, 2018). In this short communication, we report on the use of ^13^C-NMR to examine how different *Lactobacillus* species (*L. crispatus* and *L. jensenii*) differentially metabolise glucose, L- and/or D-lactate to propagate their survival and dominance.

## Material and Methods

### Bacterial culture

One strain each of two common vaginal *Lactobacillus* species *L. crispatus* (ATCC 33820) and *L. jensenii* (ATCC 25258) were cultured in De Man, Rogosa and Sharpe broth (MRS, Oxoid CM0359, Thermo Scientific, Bassingstoke, UK), under anaerobic (80% N_2_, 10% CO_2_, 10% H_2_) condition at 37°C for 24h. After this time, 50 μl of bacteria in broth was transferred to 400 μl of fresh broth and then subcultured with the addition of 50 μl 100 mM ^13^C_u_-glucose, ^13^C_3_-D-lactate, ^13^C_3_-L-lactate (or 25 μl each when ^13^C_u_-glucose and ^13^C_3_-D/L-lactate were combined) for a further 24 hours (all sourced from Sigma Aldrich, Gillingham, UK). Six separate incubation were performed for each species. Post incubation, samples were stored at −80°C until ^13^C-NMR scanning. Broth only (non-inoculated medium) and bacterial samples without ^13^C-labelled substrates were used as controls. The amount of bacterial colony forming units (CFU)/μl were estimated at time of ^13^C-substrate addition using a Helber counting chamber with Thoma ruling (area = 1/400 mm^2^, depth = 0.02 mm; Hawksley Z30000).

### NMR acquisition and processing

^13^C-NMR samples were prepared containing 430 μl bacterial sample in a 5 mm NMR tube (Norell, Morganton, NC, USA) with 20 μl D_2_O (Sigma Aldrich) and 10 μl of 200 mM ^13^C-urea (chemical shift and concentration reference, Sigma Aldrich) and 15 μl of penicillin/streptomycin (tube concentration ~105 units/ml penicillin and ~105 μg/ml streptomycin, Sigma Aldrich).

Spectra were acquired by a 9.4T Bruker AVANCE III NMR spectrometer (Bruker BioSpin GmbH, Karlsruhe, Germany) using a ^13^C{^1^H} inverse-gated pulse sequence (Spectral Width = 239 ppm, Number of acquisitions = 4096, Acquisition Time = 0.5 s, Delay Time = 2 s, Time domain points = 24036, flip angle = 16°) at room temperature (21°C). Each acquired spectrum was apodised with a 5 Hz exponential line broadening function, phase and baseline corrected using Bruker Topspin v3.4 software and referenced to the urea signal at a frequency offset δ = 165.5 ppm.

### Metabolite integration and normalisation

Spectrum integrals were determined using a custom peak fitting algorithm (Matlab R2018b, Mathworks, Natick, MA, USA) to identify and separate overlapping peaks where singlet and doublet peaks were present at a particular chemical shift (Fig. 1a: ^13^C_3_-lactate, 22.1 – 23.05 ppm; ^13^C_3_-acetate, 23.2 – 26.5 ppm). Doublet peaks were summed to give an integral for lactate or acetate formed from ^13^C_u_-glucose metabolism (Fig. 1b).

**Figure 1:**
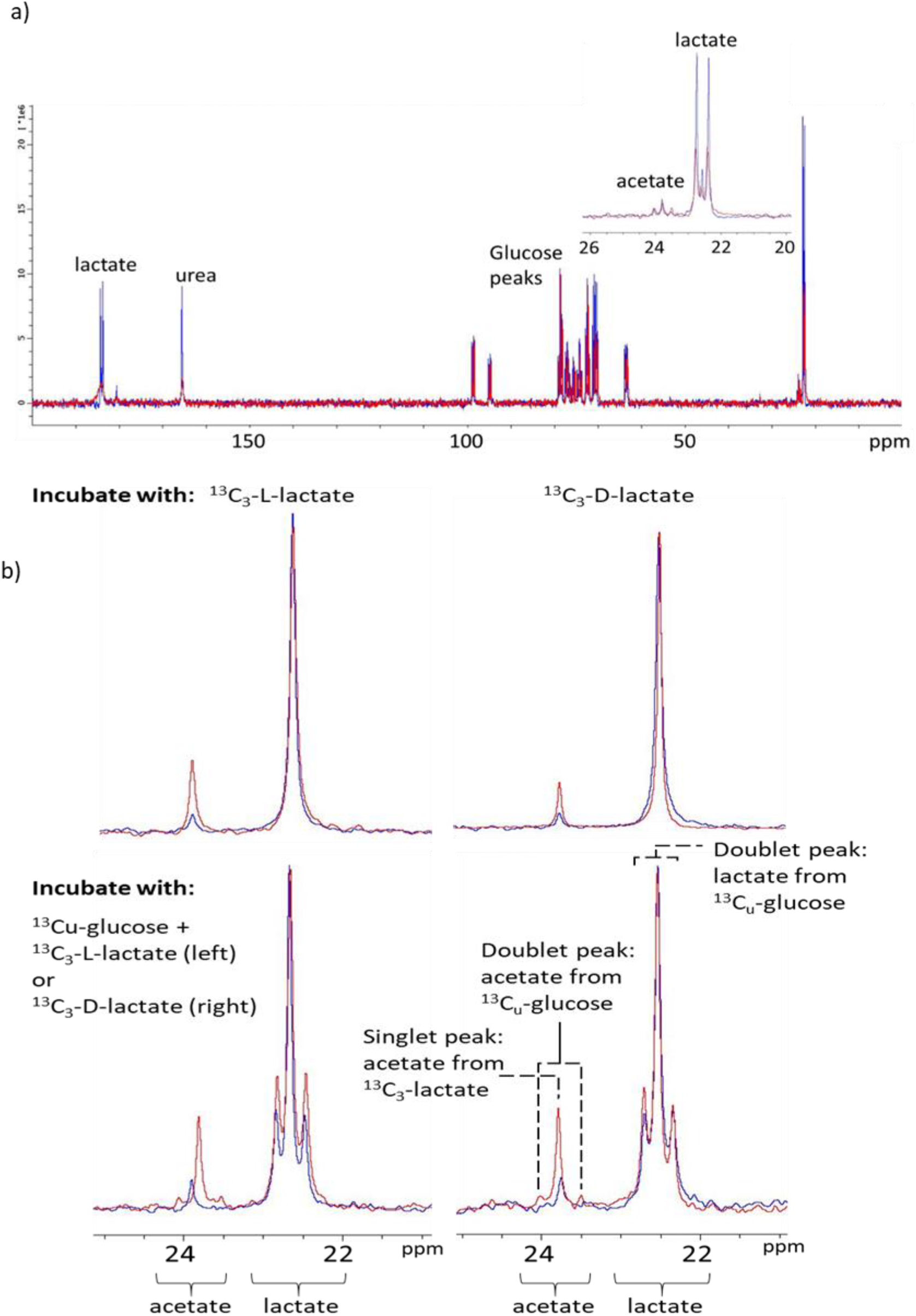
**a)** Overlaid ^13^C-NMR spectra of *L. crispatus* (blue) and *L. jensenii* (red) incubated with ^13^C_u_-glucose. Inset shows a zoomed region of the spectrum 20 – 26 ppm. **b)** *L. crispatus* (blue) and *L. jensenii* (red) incubated with either: top, ^13^C_3_-L-lactate (left) or ^13^C_3_-D-lactate (right); bottom, ^13^C_u_-glucose + ^13^C_3_-L-lactate (left) or ^13^C_u_-glucose + ^13^C_3_-D-lactate (right). Spectra scaled to lactate peak height. Doublet peaks of lactate and acetate arise from the conversion of ^13^C_u_-glucose. Singlet peaks arise from either singly labelled ^13^C_3_-D/L-lactate or substrates with natural abundance ^13^C (1.1%).

Metabolite integrals were normalised by the concentration of live bacteria present in the incubated samples. This was estimated by measuring the lactate integral, Lac_Broth_, from the conversion of broth glucose (i.e. not ^13^C-labelled) after 48 hours of incubation (24h prior to ^13^C-substrate addition plus a further 24h incubation post ^13^C addition) and was not observed in broth only samples. For each incubation experiment all integrals were divided by Lac_Broth_.

### Statistical analysis

Statistical analysis was also performed using Matlab with D’Agostino-Pearson’s tests for normality of data and a Kruskal-Wallis with Bonferroni post-hoc test (KW-B) for comparison of multiple groups with p < 0.05 taken as significant. Comparison between lactobacilli acetate and lactate integrals were performed using either Student’s t-test or Wilcoxon rank-sum test depending on the outcome of the normality test.

## Results and Discussion

The average count (CFU) of each bacterial species before incubation with ^13^C-labelled substrates was: *L. crispatus =* 4.4×10^7^ ± 7.0×10^6^ CFU/μl; and *L. jensenii* = 7.3×10^7^ ± 1.1×10^7^ CFU/μl (p = 0.1). The ^13^C-spectra showed conversion of glucose (natural abundance and ^13^C-labelled) to lactate (22.8 ppm) and acetate (23.8 ppm) for both species (n = 6 per species, Fig. 1). Higher quantities of lactate were produced by *L. jensenii* than *L. crispatus* for all ^13^C_u_-glucose containing incubations although these were not significant (Fig. 2a). The addition of ^13^C_3_–L/D-lactate to the incubation suppressed the conversion of ^13^C_u_-glucose to lactate (Fig, 2a). *L. jensenii* showed significantly higher conversion of both enantiomers of ^13^C_3_-lactate to acetate (23.8 ppm) than *L. crispatus* (*L. crispatus* vs *L. jensenii*: D-lactate to acetate: 0.26 ± 0.10 vs 1.71 ± 0.15, p < 0.001; L-lactate to acetate: 0.11 ± 0.04 vs 1.68 ± 0.14, p < 0.001, Fig. 2b). Six of the *L. jensenii* ^13^C spectra (^13^C-glucose, n = 2; ^13^C-glucose/L-lactate, n = 2; ^13^C-glucose/D-lactate, n = 1; ^13^C-L-lactate, n = 1) also showed peaks at 32.4 and 180.6 ppm (assigned as succinate from HMBC spectra, data not shown).

**Figure 2:**
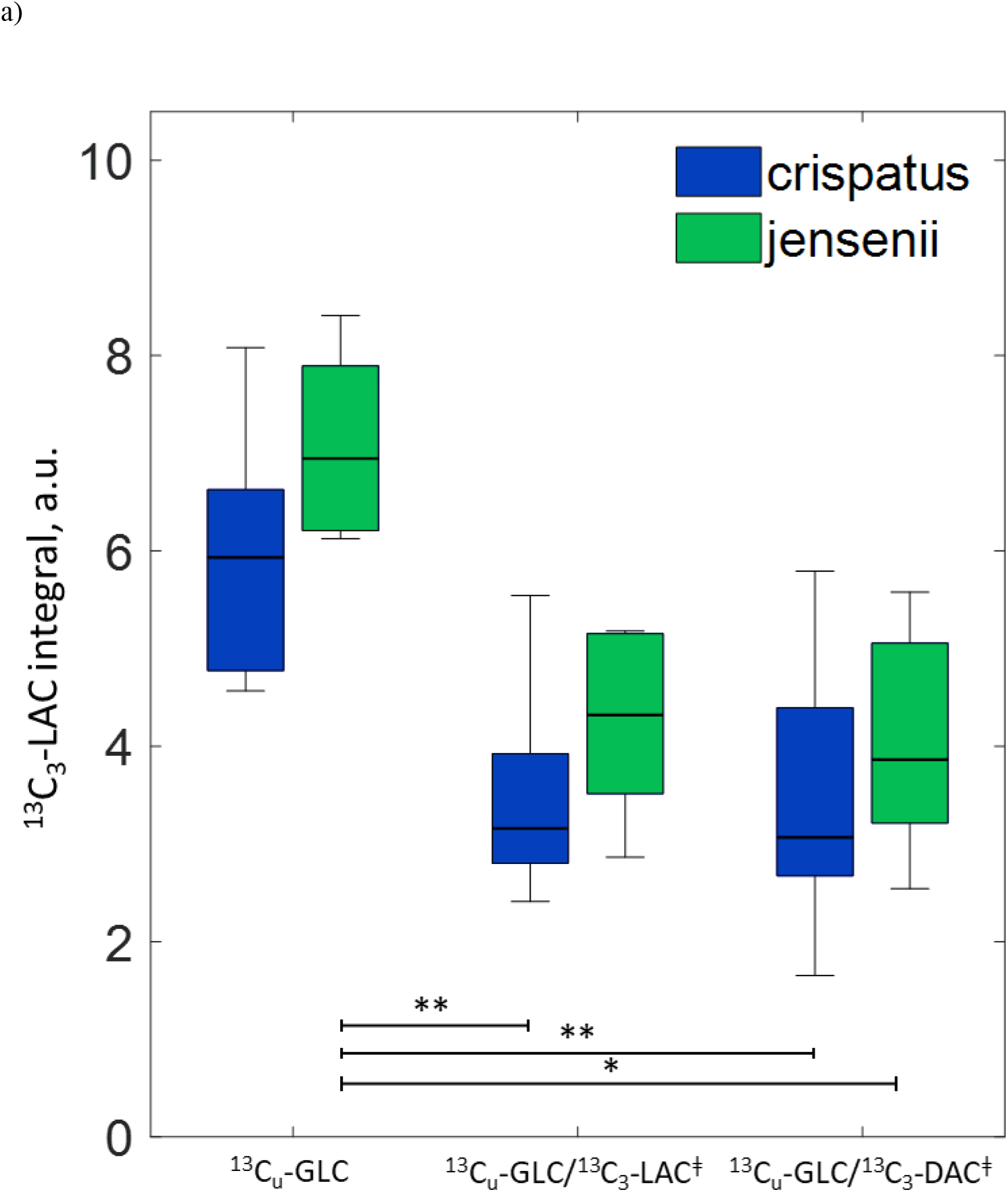
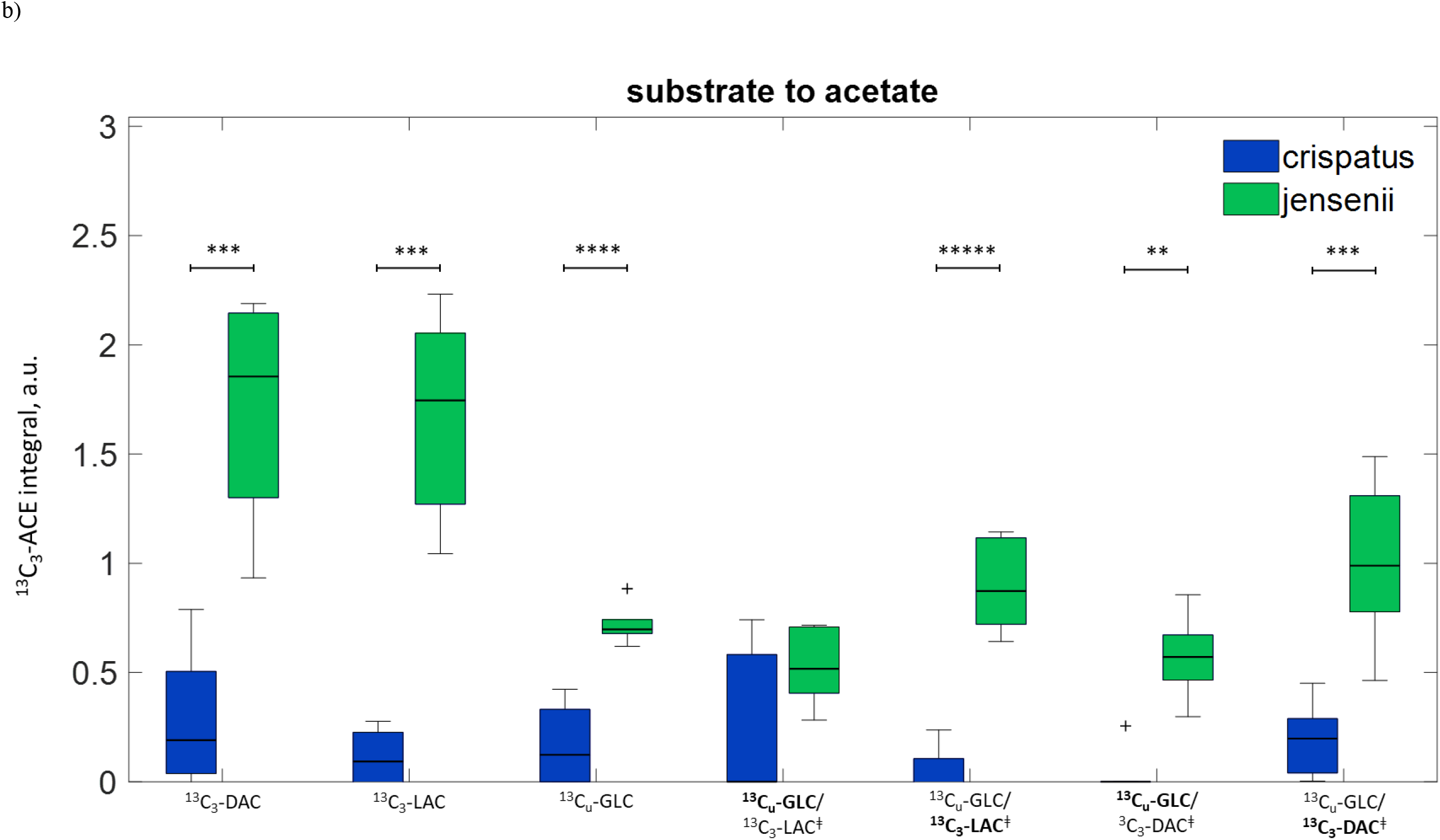
**a)** Normalised ^13^C_3_-lactate integrals for the conversion of ^13^C_u_-glucose (GLC), ^13^C_u_-glucose + ^13^C_3_-L-lactate (LAC) or ^13^C_u_-glucose + ^13^C_3_-D-lactate (DAC). ^‡^ For combination incubations, the source substrate that is converted to lactate is ^13^C_u_-glucose (as detected from doublet peak in spectrum). Kruskal-Wallis with Bonferroni post-hoc test: *p < 0.05, **p < 0.01. **b)** Normalised ^13^C_3_-acetate (ACE) integrations for the conversion of ^13^C_u_-glucose (GLC), ^13^C_u_-glucose + ^13^C_3_-L-lactate (LAC) or ^13^C_u_-glucose + ^13^C_3_-D-lactate (DAC). ^‡^ For combination incubations, the source substrate that is converted to acetate is highlighted in bold. Boxplots show median, interquartile range (IQR) and whiskers (1.5xIQR). Wilcoxon rank sum: ** p < 0.01, *** p < 0.001, **** p < 0.0001, ***** p < 0.00001.

Results show that *Lactobacillus* species that are present in the vagina differentially metabolise ^13^C-labelled glucose and lactate to produce acetate and other metabolites. *L. jensenii* spectra showed that this species was able to convert more D- or L-lactate to acetate than *L. crispatus*. D-lactate is believed to be a more potent anti-infection and anti-inflammatory agent than L-lactate (van de Wijgert *et al*., 2014; Witkin and Linhares, 2017; Witkin *et al*., 2013) and other studies have shown that *L. crispatus* produces more D-lactate than *L. jensenii* (Amabebe and Anumba, 2018; van de Wijgert *et al*., 2014; Witkin and Linhares, 2017; Witkin *et al*., 2013). This study observed that *L. jensenii* converted a higher amount of ^13^C_u_-glucose to lactate than *L. crispatus*. We did not ascertain if the lactate produced was either the L- or D isomer. However, this could be done by either an NMR chiral shift reagent (Zhang *et al*., 2017a; Zhang *et al*., 2017b) or absolute quantification of these isomers by enzyme-based spectrophotometry as we have previously demonstrated (Amabebe *et al*., 2019; Amabebe *et al*., 2016a; Cavanagh *et al*., 2019).

Bacterial load (CFU) was not used for normalisation, as this did not correlate with ^13^C-lactate integrals observed in the spectra (data not shown). Hence, the integrals in this study were normalised based on the assumption that the observed lactate peak arising from non-^13^C-labelled glucose present in the media (singlet peak between the doublet for lactate at 22.8 ppm, Fig 2a) was proportional to the concentration of live bacteria. Justification for this approach was made by the observation of significant correlations between the doublet peak ^13^C-lactate integrals, (^13^C-glucose to lactate metabolism), and singlet peak lactate integrals (broth glucose to lactate. Data not shown). Using this approach could potentially underestimate the quantity of live bacteria due to further metabolism of lactate, e.g. to acetate. Considering acetate concentration is confounded by its presence in the broth, which then overestimates the bacteria concentration (acetate integrals from broth only spectra would need to be measured to account for this). Using CFU for normalisation requires careful assessment of the inhomogeneity in bacteria distribution that can make this method inaccurate. Additionally, the viability of bacteria would need to be determined, as some of the bacteria will be dead. Furthermore, bacteria that were metabolically active at the start of the incubation would die over the duration of the incubation leading to an over correction of the integrals.

The conversion of glucose and lactate to acetate and succinate by *L. jensenii* compared to *L. crispatus*, suggests that this may be an important pathomechanism of dysbiosis, altered vaginal pH, infection and infection-associated spontaneous preterm birth. Elevated amounts of acetate and succinate in the vaginal space is associated with increased pH and risk of infection such as bacterial vaginosis (Aldunate *et al*., 2015; Ceccarani *et al*., 2019), a known risk factor of preterm birth and other poor reproductive outcomes (Amabebe and Anumba, 2018). Vaginal bacterial communities dominated by *L. jensenii* usually have higher pH compared to when *L. crispatus* predominates (Aldunate *et al*., 2015). We have previously observed that symptomatic women at risk of preterm birth show elevated cervicovaginal fluid acetate (Amabebe *et al*., 2016a; Amabebe *et al*., 2016b), which can be combined with pro-inflammatory mediators to improve its performance as a predictive marker (Amabebe *et al*., 2019).

These experiments were performed on bacteria that had been freeze thawed. As NMR is a non-destructive technique, it would be possible to apply this method to live bacteria to sequentially acquire spectra and determine the rate of conversion of substrates. Whilst it would be important to characterise the most common vaginal strains of these species, alternative strains could also be characterised. Future studies will be expanded to other *Lactobacillus* species and strains such as *L. iners* and *L. gasseri* and anaerobes including *Gardnerella*, *Bacteroides* and *Mobiluncus*, as well as fungi (e.g. *Candida albicans*) that are known to exist within the vaginal milieu.

## Declarations

### Funding

This research did not receive any specific grant from funding agencies in the public, commercial, or not-for-profit sectors. However, EA and DA are funded by National Institute for Health Research (NIHR, 17/63/26).

### Conflict of interest statement

The authors have no conflicts of interest to declare.

### Author contributions

This work was conceived and conceptualised by EA and SR. EA and SR acquired and analysed the data with DA supervising. All authors contributed to drafting and review of the manuscript, and approved the final version for submission.

## Acknowledgements

We are grateful to the University of Sheffield, UK, for providing support and the 9.4T MRS spectrometer with which this study was performed.

